# Repurposed high-throughput images enable biological activity prediction for drug discovery

**DOI:** 10.1101/108399

**Authors:** Jaak Simm, Günter Klambauer, Adam Arany, Marvin Steijaert, Jörg Kurt Wegner, Emmanuel Gustin, Vladimir Chupakhin, Yolanda T. Chong, Jorge Vialard, Peter Buijnsters, Ingrid Velter, Alexander Vapirev, Shantanu Singh, Anne Carpenter, Roel Wuyts, Sepp Hochreiter, Yves Moreau, Hugo Ceulemans

## Abstract

We repurpose a High-Throughput (cell) Imaging (HTI) screen of a glucocorticoid receptor assay to predict target protein activity in multiple other seemingly unrelated assays. In two ongoing drug discovery projects, our repurposing approach increased hit rates by 60- to 250-fold over that of the primary project assays while increasing the chemical structure diversity of the hits. Our results suggest that data from available HTI screens are a rich source of information that can be reused to empower drug discovery efforts.

---

High-throughput (cell) Imaging (HTI) captures the morphology of the cell and its organelles by high-throughput microscopy and is successfully applied in many areas of current biological research(1–3). In a pharmacological context, HTI is applied to screen chemical compounds based on morphological changes they induce(4, 5). Currently, most HTI screens are designed for the single purpose of evaluating one specific biological process and exploit only a handful of morphological features from the image(6), see Fig. 1b. These morphological features are understood to directly reflect that biological process. However, any imaged cellular system hosts many more biochemical processes and thousands of potential drug targets, all of which are exposed to the screened chemical compounds. Many of these targets and processes impact cell morphology and that morphology can to a large extent be extracted from the images(7). This set of features can be used to describe chemical compounds and can be considered as an image-based fingerprint. Wawer et al.(8) proposed the use of image-based fingerprints to optimize the diversity of medium scale compound sets. Image-based fingerprints can also be used to group compounds by pharmacological mechanism(9). Thus, images provide a rich source of biological information that can be leveraged for a variety of purposes in drug discovery.

**Fig. 1.**
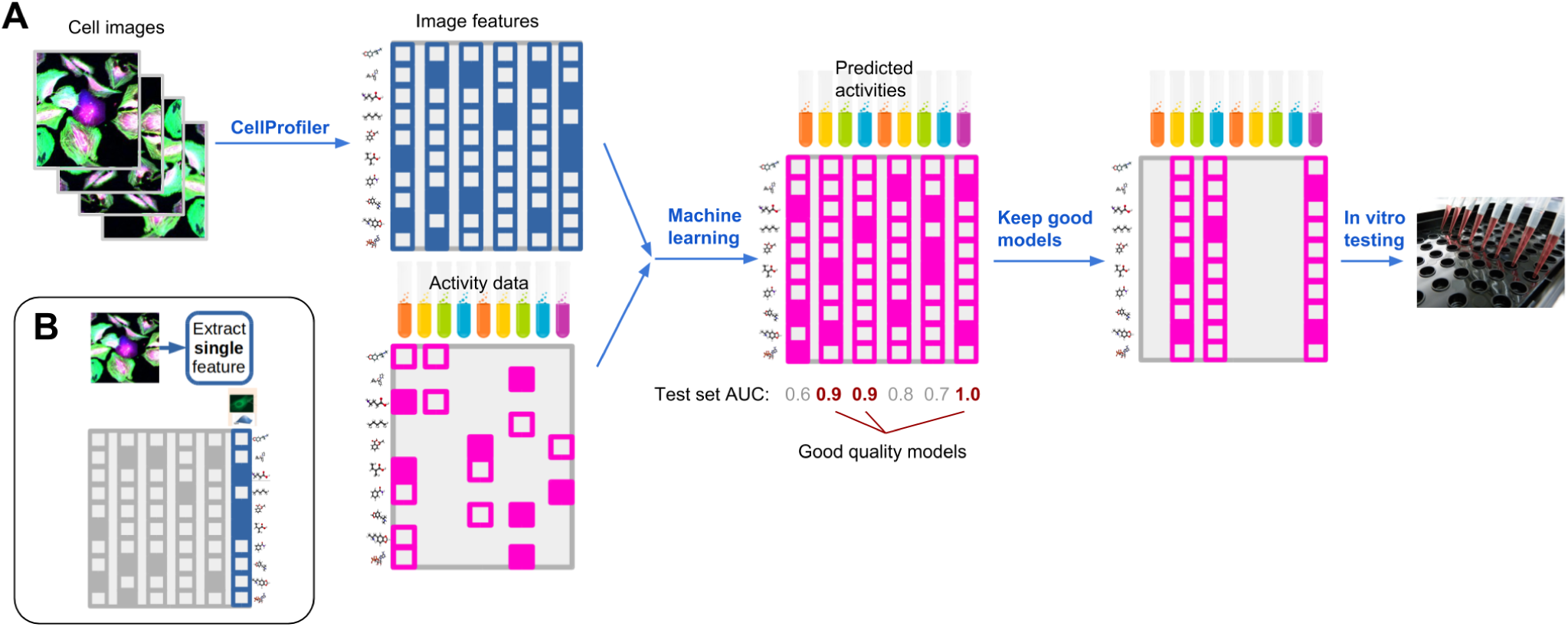
Repurposing imaging screens. Panel A: Our repurposing approach is depicted. A large number of features are extracted from images of cells which are then used by machine learning methods to model all available activity data from previously performed assays. Targets with good predictivity on the test data are then selected for in vitro validation. Panel B: A typical HTI screen approach is depicted (10, 11). Few or single features are extracted from cellular images.

Here, we propose the systematic evaluation of image-based fingerprints from HTI screens for predictivity on a large collection of protein targets, most of which were not considered during the design of the screen (Fig. 1a). To this end, an extensive fingerprint of morphological features was extracted for each compound imaged in a single screen, aiming for maximal and unbiased information capture. We then deployed a machine learning approach to predict the activity across a broad set of validated target assays (from now referred to as assay) based on the image-based fingerprint of the compounds, and evaluated model performance for each assay. In this way, we hypothesized existing HTI screens can be repurposed to inform on the activity of untested compounds in assays for which a high-quality model is available (Fig. 1a).

This procedure is reminiscent of the predictive modeling approaches used for virtual screening and QSAR, but differs in that it uses image-based fingerprints rather than chemical fingerprints that encode the structure of compounds. Chemistry-based models are predictively performant, but only for those parts of chemical space for which sufficient assay activity data is available, because chemical fingerprints themselves do not incorporate biological or target information. Hence, compounds that are chemically very different from any known active compound are unlikely to be predicted as active. Image-based models are expected to be less dependent on the availability of chemically similar training examples. The model then correlates all imaged biology to the biological activities to predict. Therefore, image-based models could outperform chemistry-based models in novel and activity-wise poorly annotated chemical space.

In our study, we repurposed a high-throughput imaging screen of 524,371 compounds originally used for the detection of glucocorticoid receptor (GCR) nuclear translocation. Each compound was applied in a concentration of 10µM to H4 brain neuroglioma cells, incubated for one hour, before the addition of 1µM hydrocortisone to stimulate translocation of the GCR. After an additional 1 hour of incubation, cells were fixed and imaged in 3-channel fluorescence, with a nuclear stain (Hoechst), CellMask Deep Red (Invitrogen) to delineate cell boundaries, and indirect immunofluorescence detection of GCR. For repurposing the screen, the images were post-processed using CellProfiler software. Using a pipeline similar to that of Gustafsdottir and colleagues(12), we extracted unbiased maximally-informative features from the images. For each cell in the image, the pipeline computed an image-based fingerprint of 842 features. In our machine learning models, each compound was then represented by the vector of feature medians across all cells in the image.

We then built a machine-learning model using Bayesian matrix factorization with the image-based fingerprint as side information (Online Methods). This model was evaluated for its predictivity across assays that map to more than 600 drug targets leveraging more than ten million activity measurements. We assessed the performance of this model using nested cluster cross-validation(13). The model was predictive for 37.3% of the assays (AUC > 0.7 on 225 assays, Online Methods) and offered high predictivity for 5.6% of the assays (AUC > 0.9 on 34 assays, Online Methods). Among these 34 assays with high-quality predictions, two were running in ongoing discovery projects: one oncology project and one central nervous system (CNS) project. We used the matrix factorization model to select compounds for testing by these two projects.

For the oncology project, the target was a kinase with no known direct relation to the glucocorticoid receptor. Using our matrix factorization model, we ranked about 60,000 compounds tested in the GR assay but for which no activity measurement was available in the oncology screen. We then experimentally validated the 342 compounds ranked highest by our matrix factorization method, roughly the amount of non-control wells on a plate. This resulted in 141 submicromolar hits (41% hit rate), which corresponds to a 60-fold enrichment over the initial HTS (0.725% hit rate).

To evaluate the chemical diversity of the hits, we computed the Tanimoto similarity (based on extended-connectivity fingerprints (ECFP)(14)) of each hit to the nearest hit identified by the initial HTS. When compared to that of the initial hits (red distribution in Fig. 2a), the distribution of these similarities implies a substantially improved chemical structure diversity (shift to the left) of the image-based hits (green). As a reference, the figure also depicts the distribution for randomly selected compounds (blue). Thus, the HTI matrix factorization model selected candidate compounds with a high hit rate, and diversified the hit space.

**Fig. 2.**
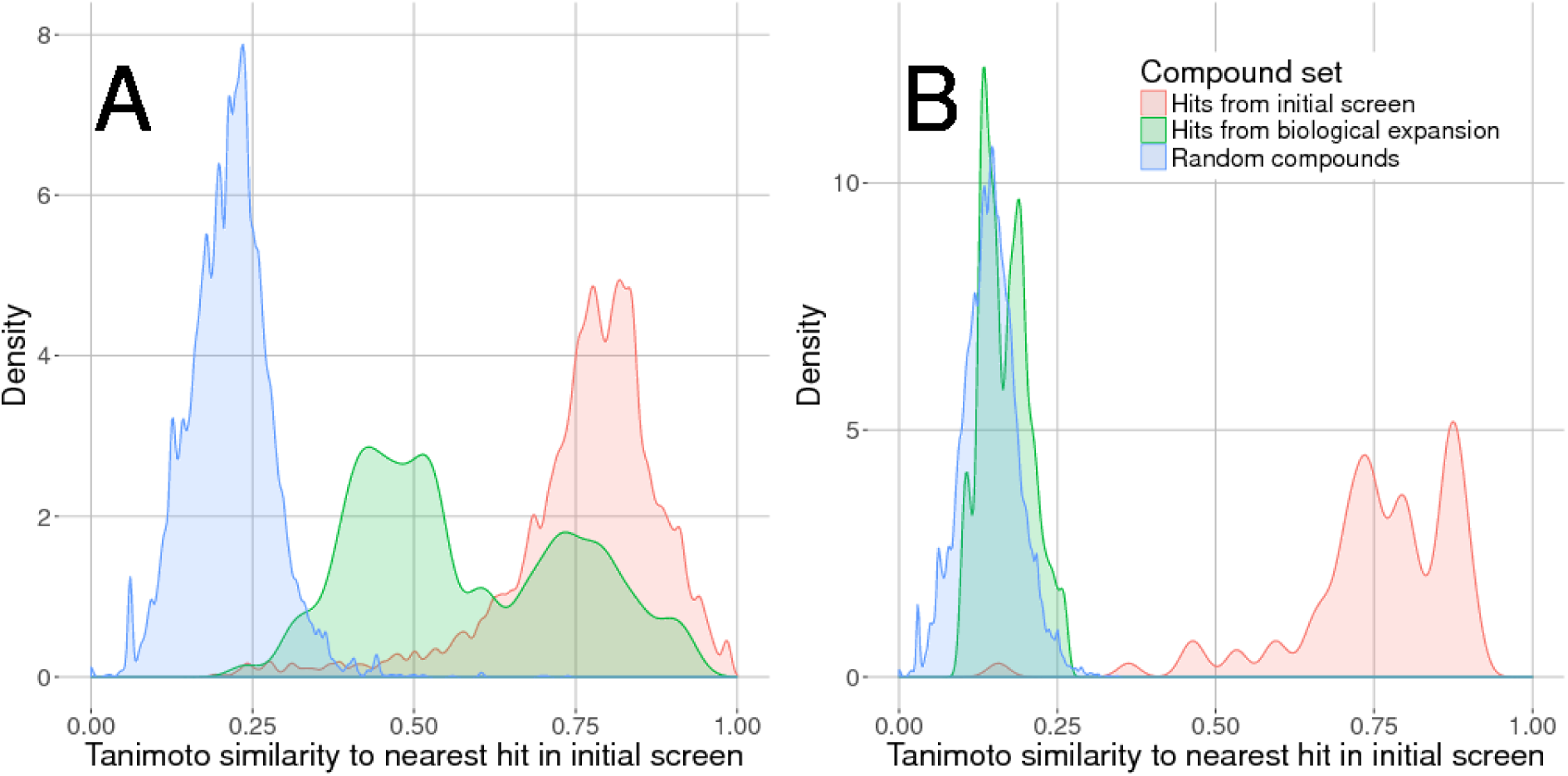
For each compound in a set, the ECFP (radius 4) based Tanimoto similarity to the nearest hit from the initial screen was considered. Similarity densities for the set of initial hits, the set of hits from our biological extension and a randomly selected set of compounds are depicted in red, green and blue, respectively, for the oncology project in Panel A and for the CNS project in Panel B. Note that in the CNS project, unlike the oncology project, the selection procedure involved an additional step to reduce representatives from the same chemical-structural class. Overall, the hits discovered by our approach are chemically highly diverse.

For the CNS project, the target was a non-kinase enzyme again without obvious relation to the glucocorticoid receptor. Using our matrix factorization model, activity was predicted for all 500,000 image-annotated compounds. Compounds with predicted submicromolar activity were filtered to deplete for unfavorable properties(15), like autofluorescence and low predicted central nervous system availability (Online Methods), and the remaining compounds were grouped in chemical clusters from which we randomly selected a handful of representatives from each cluster (Online Methods). The 141 compounds resulting from this procedure were experimentally tested, and for 37 of them, submicromolar activity was confirmed, resulting in a 22.7% hit rate or a 250-fold enrichment over the hit rate of the initial HTS (0.088%). Importantly, the 37 hits mapped to 32 Murcko scaffolds(16) that were not represented in the set of initial hits. The distribution of Tanimoto similarities to the nearest hit in the initial screen (Fig. 2b) supports that conclusion. These results again suggested that an image-empowered compound selection strategy can boost hit rate and hit diversity.

To check whether the success of our approach arises from the machine learning method or from the description of chemical compounds by imaging features, we applied three different machine learning methods. We used Macau(17), a regression method based on Bayesian matrix factorization with side information (Online Methods), random forest classification(18) and deep neural networks(13) (Online Methods). These machine learning models performed similarly in terms of the assays that could be predicted accurately (Supplementary Fig. S4 and S5). Our cross-validation setup also guarantees that the predictive performance does not come from the activity data of the same compound across other targets (Online Methods). Therefore, we conclude that the description of compounds by imaging features is the essential contribution to the success of our approach.

We emphasize that the method is a supervised machine learning method and hence output labels (in this context, activity measurements) are required to train the model. This requires that activity measurements be acquired for a reasonably sized library of compounds.

Our results indicate that images from HTI screening projects that are conducted in many institutions can be re-purposed for increasing hit rates in other projects, even those that seem unrelated to the primary purpose of the HTI screen. Consequently, it might be possible to replace particular assays with the potentially more cost-efficient imaging technology together with machine learning models. By accessing rich morphological features of the cell, imaging screens provide a view over various cellular processes resulting in a fingerprint of biological action. This raises an interesting question of the breadth of targets that could be accessed by imaging screens if different cell lines, culturing conditions, staining of organelles and/or incubation times are used.

The focus of this report was to demonstrate that the use of HTI data enables the identification of diverse hits without the need to retest the entire library in the target assay. We note that our models may also support target deconvolution for phenotypic screens, through the prioritization of targets with predicted activities that match phenotypic observations.

Moreover, in the light of recent advances of convolutional neural networks, raw images might be used directly in the activity prediction pipeline. This would allow the model to learn the best image features for the specific task at hand and may strengthen our approach. Furthermore, our results are based on a single HTI screen and we envision that a collection of multiple HTI screens could even be more powerful with respect to assay activity prediction. Finally, our imaging features are median values across all cells from an image. However, models based on the distribution of the feature values (e.g. quantiles) or even single cell analysis could prove to have higher predictive power and will be investigated in the near future.

## ACKNOWLEDGMENTS

This work was supported by research grants IWT135122 ChemBioBridge, IWT130405 ExaScience Life Pharma and IWT150865 Exaptation from the Flanders Innovation and Entrepreneurship agency. The NVIDIA Corporation generously donated a GPU. J.S., A.A., and Y.M. were additionally supported by Research Council KU Leuven: CoE PFV/10/016 SymBioSys, PhD grants and imec strategic funding 2017.

## Author contributions

J.S., G.K., A.A., S.H., Y.M., and H.C. conceived this study and designed the experiments. J.S., G.K., A.A., M.S., J.K.W, E.G., V.C., Y.T.C., J.V., P.B., I.V., A.V., S.S., A.C., R.W., and H.C. conducted the experiments. S.H., Y.M., and H.C. supervised this project. J.S., G.K., A.A. and H.C. wrote the manuscript with input from all authors.

## Online Methods

In this Online Methods, we provide details on the data set, provide a description of the used machine learning methods and the method comparison and evaluation.

## Data set

The data set comprises 524,371 samples, 842 features and 1,200 prediction tasks. The samples correspond to chemical compounds that were administered to cells. The features are derived from a biological imaging technique together with the CellProfiler software that calculates morphological features of the imaged cells. This means that a chemical compound is characterized by its induced morphological changes of cells. The prediction tasks comprised compound activities that were measured by multiple independent biological experiments, so-called assays. Typically, a single compound is measured in one or very few assays, such that the matrix that has to be predicted is populated by many unknown (“NA”). values. In total there are more than 10 million activity values for the 1,200 prediction tasks, resulting in fill rate of about 1.6%. The samples were distributed across three cross-validation folds to enable the nested cluster cross-validation [1] (see Section “Cross-validation and measuring predictivity”).

## Macau: Bayesian matrix factorization

We built a Bayesian matrix factorization method called Macau that incorporates image features from CellProfiler as side information. To factorize the 524 371 × 1 200 activity matrix **Y**, Macau represents each compound and each assay by *D*-dimensional latent vectors **u**_i_ and **v**_j_, respectively. The prediction for the element *Y_ij_*, corresponding to the activity of compound *i* on assay *j* is given by the scalar product 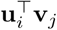. The image features x*_i_* are 842 dimensional vectors (see Data set section) and are added to the prior of the latent vectors of compounds u*_i_*.

This results in a probabilistic model of 
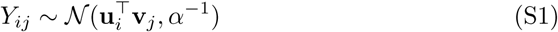

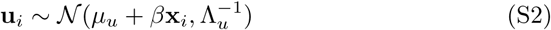

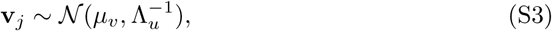
where *α* is precision of the observations, parameters *µ*_u_ and Λ_u_ model the mean and precision of the compound latent vectors, similarly *µ_v_* and Λ*_v_* model the latent vectors for assays. The parameter *β* is a D × 841 dimensional matrix that maps the image features to the compound latent space. To learn *β* we apply Gaussian prior on it: 
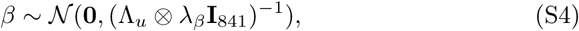
where ⊗ is the Kronecker product, λ*_β_* is precision parameter and **I**_841_ is the identity matrix of size 841. The Figure S1 depicts the plate diagram for the probabilistic model.

By deriving conditional probabilities we obtain Gibbs sampler for the model, as was done in [2]. We use Gibbs sampler to learn the parameters of the model, except for *α* and *D* that are fixed to 10 and 150, respectively. For the experiments, we ran Gibbs sampler for 2000 iterations and discarded the first 400 as burn-in. To compute the final answer we use each Gibbs sample to make a prediction for *Y_ij_* and then we compute the average over the samples.

To evaluate the model performance we used 3-fold cross-validation as described in the next section.

## Cross-validation and measuring predictivity

In our *in silico* experiments we used 3-fold cross-validation where all of the compounds of a chemical series are in a single fold (see Figure S2). Each chemical series was defined by 0.7 Tanimoto similarity cutoff on ECFP6 features. Therefore, the test set never contains a compound that is chemically very close to a training set compound. With this setup we can measure the model’s actual ability to generalize across series.

For each cross-validation fold we compute its test AUC-ROC scores for each target assay at pIC_50_ thresholds of 5.5, 6.5, 7.5 and 8.5. We use the average of the 3 folds as the evaluation metric for each assay-threshold pair. We then only keep the assay-threshold pairs with at least 25 actives and 25 inactives, and consider the model to have high (moderate) predictivity on an assay if one of the thresholds has AUC-ROC higher than 0.9 (0.7).

**Figure S1.**
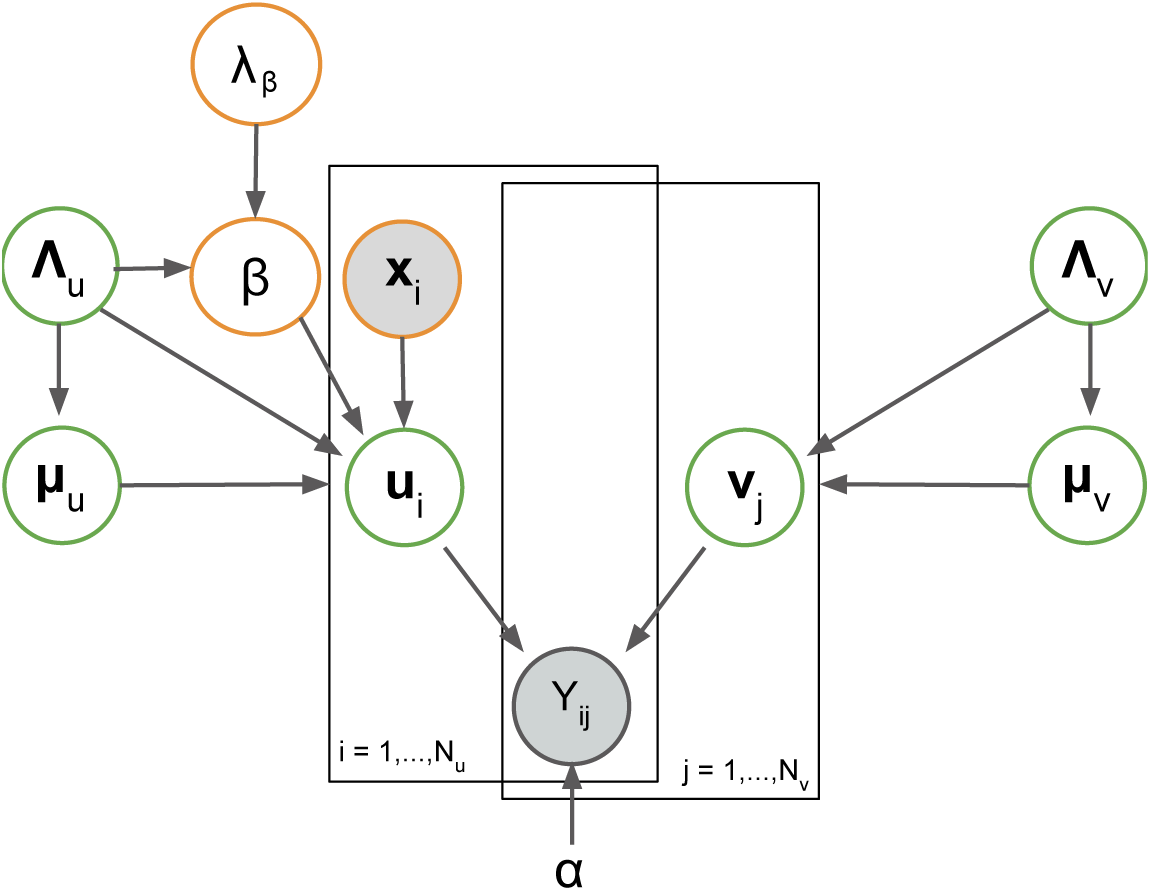
Diagram for the probabilistic model for the used matrix factorization approach Macau.

**Figure S2.**
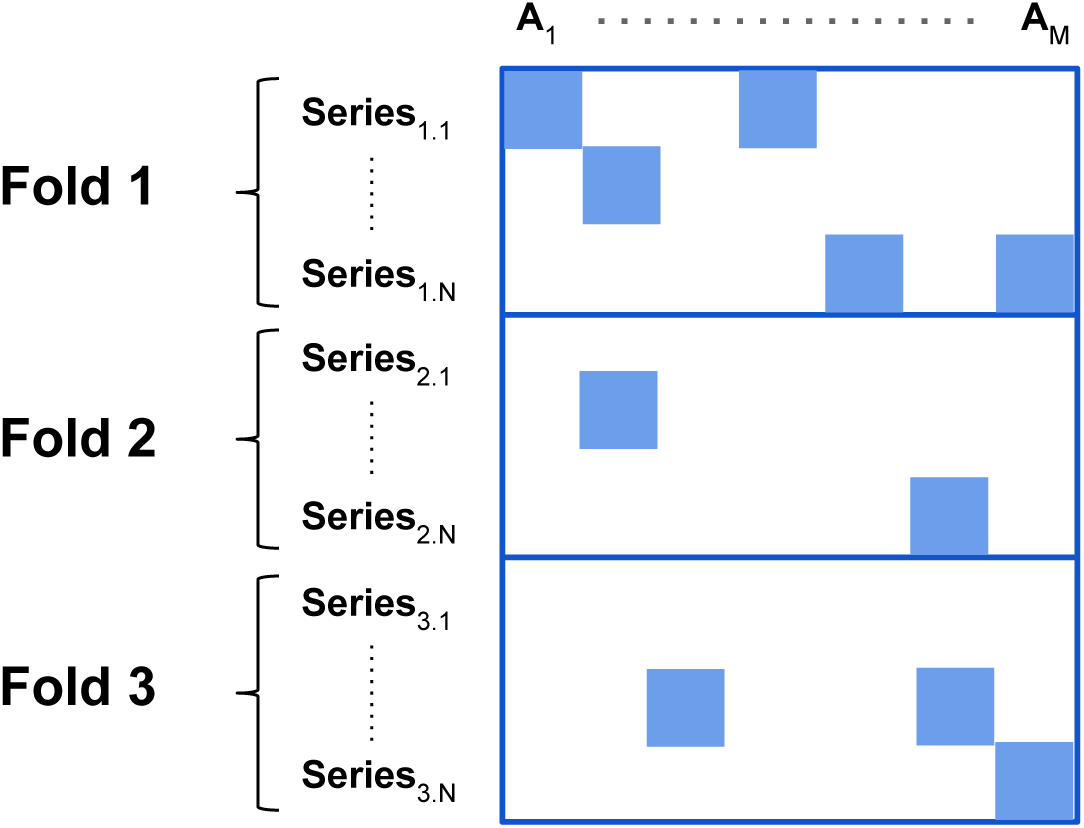
Clustered cross-validation for compound-protein activity prediction. Every block of rows corresponds to a compound series and every column to anassay.

## Autofluorescence filtering and CNS availability

Frequent hitters are compounds that are promiscuously active, e.g. based on certain substructure motifs they might contain. Also, some compounds might be dyes themselves, be reactive species, or interfere with particular assay technologies as Fluorescent or AlphaScreen readouts. Baell/Holloway [3] suggested a Pan Assay Interference Compounds (PAINS) filter for removing such promiscuous compounds from HTS hits.

The Blood-Brain-Barrier (BBB) is a critical membrane to separate the blood from the brain in the central nervous system (CNS). Drugs for CNS disease indications should pass the BBB, while drugs for non-CNS indications should not pass the BBB for preventing unwanted side-effects. The BBB allows the passage of water and lipid-soluble molecules by passive diffusion. Two major estimations for BBB permeability are therefore based on passive diffusion models based on logP and polar surface area (PSA) of compounds [4], or active transport via a P-glycoprotein (P-gp) substrate probability of compounds [5, 6]. We filtered out all compounds that do not exhibit BBB permeability according to standard pharmaceutical practice.

## Deep neural networks (DNNs)

We implemented Deep Neural Networks (DNNs), concretely feed-forward artificial neural networks, with many layers comprising a large number of neurons [1]. DNNs consists of interconnected *neurons* that are arranged hierarchically in layers. In the first layer of the network (the *input layer*), the neurons obtain an input vector which are the descriptors of the chemical compound. The intermediate layers (the *hidden layers*) comprise the *hidden neurons* which have weighted connections to the neurons of the lower level layer, and can be considered as abstract features, built from features below. The last layer (the *output layer*) supplies the predictions of the model. While traditional networks employed a small number of neurons, a DNN may have thousands of neurons in each layer [7]. Figure S3 shows the general architecture of Deep Neural Networks.

### HCI features as network inputs

Each chemical compound is represented by image features from CellProfiler averaged across multiple cells. These 841 features entered the input layer of the DNNs.

### Activation function

We used rectified linear units (ReLUs) as activation functions in the hidden layers. The output layer has sigmoid activation functions.

### Regularization

To avoid overfitting, we employed multiple regularization techniques, concretely *Dropout* [8] and *early stopping*. Both the dropout rate and the early-stopping parameter, that is the number of epochs after which learning is stopped, were determined on a validation set.

**Figure S3:**
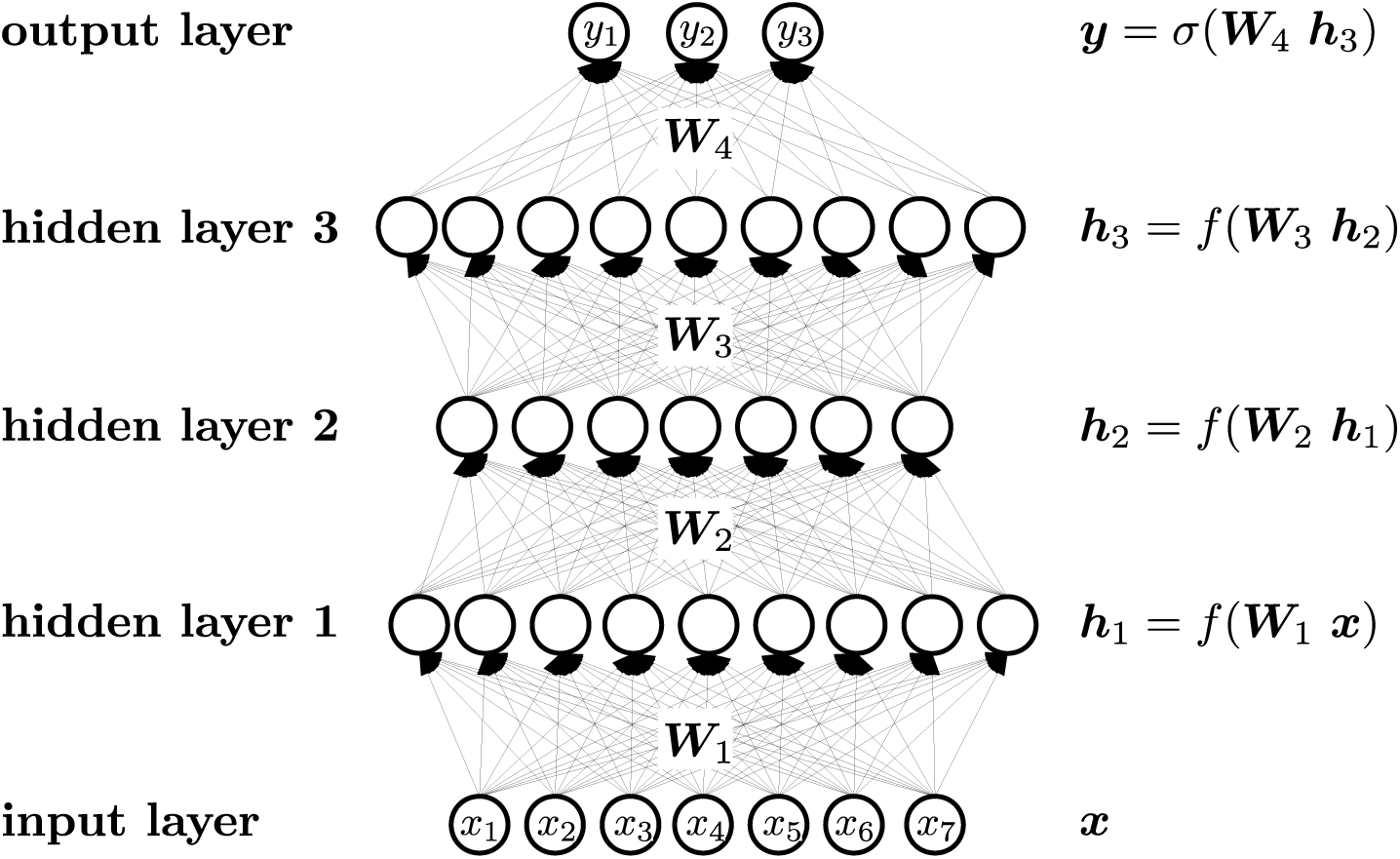
Architecture of Deep Neural Networks

### Multitask Learning

Since each chemical compound can have multiple biological activities, the prediction task at hand represents a multi-task setting. Deep Learning naturally enables multi-task learning [9]. These tasks can mutually increase the representation of the samples in the hidden layers which can further lead to performance improvements [1]. We modeled each assay by a separate output unit, which led to DNNs with around 1,200 output units.

### Objective Function

We used cross-entropy as a loss function for our DNNs:

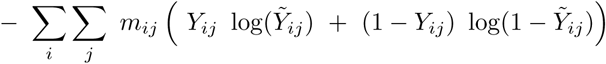
where 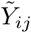 is the prediction for compound *i* and assay and the actual label is *Y_ij_*. *Y_ij_* indicates whether the compound was active (*Y_ij_* = 1) or inactive (*Y_ij_* = 0) in this assay. Additionally, for many compounds the activity has not been measured which we accommodated by a binary variable *m_ij_* that is zero if a measurement is present and zero otherwise.

### Training of the DNN and Software implementation

We used minibatch Stochastic Gradient Descent (SGD) to train the DNNs. Hence, we implemented the DNNs using the CUDA parallel computing platform and employed NVIDIA Tesla K40 GPUs to achieve speed-ups of 20-100x compared to CPU implementations.

### Hyperparameter search

We optimized the DNN architecture and hyperparameters, such as learning rate, early-stopping parameter and Dropout rate on a validation set in a nested cross-validation procedure [10, 11]. This procedure produces performance estimates that are unbiased by hyperparameter selection since the hyperparameters are optimized on the inner folds and only the best performing hyperparameters are tested on the outer folds. We considered 1, 2 or 3 layer networks with 1024, 2048, or 4096 units in each layer. The tested learning rates were 0.01, 0.05, and 0.1. The Dropout rates were either set to zero, or to 10% dropout in the input layer and 50% dropout in the hidden layers. Additionally, we tested whether the dropout rate should be arithmetically increased from 0 by 0.005 after each epoch (“dropout schedule”) until the given dropout rate or whether the dropout rates were constant (“no dropout schedule”) during learning. Table S1 summarizes these hyperparameters and architecture design parameters that were used for the DNNs, together with their search ranges.

**Table S1:** Hyperparameters considered for Deep Neural Networks

| Hyperparameter | Considered values |
| --- | --- |
| Number of Hidden Units | {1024, 2048, 4096} |
| Number of Hidden Layers | {1, 2, 3} |
| Learning Rate | {0.01, 0.05, 0.1} |
| Dropout | {no, yes (50% Hidden Dropout, 10% Input Dropout)} |
| Dropout-schedule | {no, yes} |

**Table S2:** Hyperparameters considered for Random Forests

| Hyperparameter | Considered values |
| --- | --- |
| criterion | {Gini, cross-entropy} |
| number of trees | {250} |
| number of features considered at each split | $\{1, 2, 3\} \times \sqrt{\text{total number of features}}$ |

The hyperparameters that performed best when averaged across the three cross-validation folds were: 3 layers with 2 048 units, learning rate 0.05, Dropout: yes, Dropout-schedule: yes. The early stopping-parameter was determined to be 63 epochs.

## Random Forests (RF)

Random Forests work well with different types of descriptors [12] at a large variety of tasks and their performance is relatively robust with respect to hyperparameter settings [13]. We used a high number of trees to obtain a stable model with high performance [14] The critical parameter is the number of features considered at each split [15] which we adjusted in the established nested cross-validation setting. We trained and assessed models for each assay individually in our established framework using different hyperparameters given in Table S2 and the Random Forest implementation “ranger” [16].

## Method Performance

We compared Macau, a regression method based on Bayesian matrix factorization with side information (Online Methods), random forest classification (Breiman, 2001) and deep neural networks (Mayr, 2016) (Online Methods) for predictive performance using imaging features as inputs, see Supplementary Table S1 for the detailed results. We found that the methods performed similarly with respect to which assays could be predicted accurately. Concretely, 11 assays could be predicted with AUC above 0.9 by all three methods (see Venn diagram in Figure S4). Similarly, 182 assays had performance of AUC above 0.7 by all three methods (see Venn diagram in Figure S5). The numbers of assays where only a single or a pair of methods gave an AUC above 0.7 are all comparably smaller. Therefore, we conclude that the performance is mainly driven by imaging features rather than the machine learning method.

**Figure S4:**
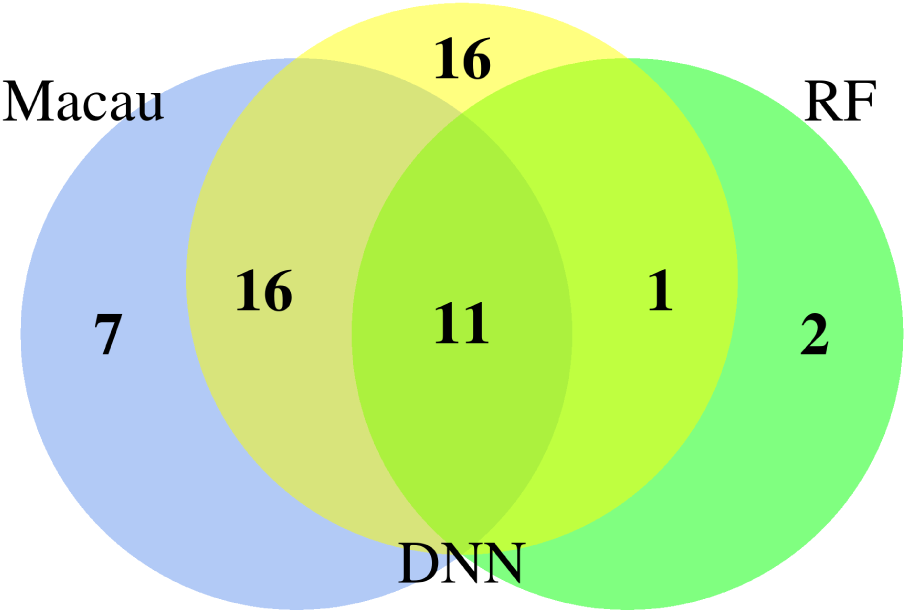
Venn diagram: number of protein targets with high predictivity (AUC > 0.9).

## External Tables

The table providing performance values for each method at each prediction task is attached as an external file: `Supplementary Table S1.xls`

**Figure S5.**
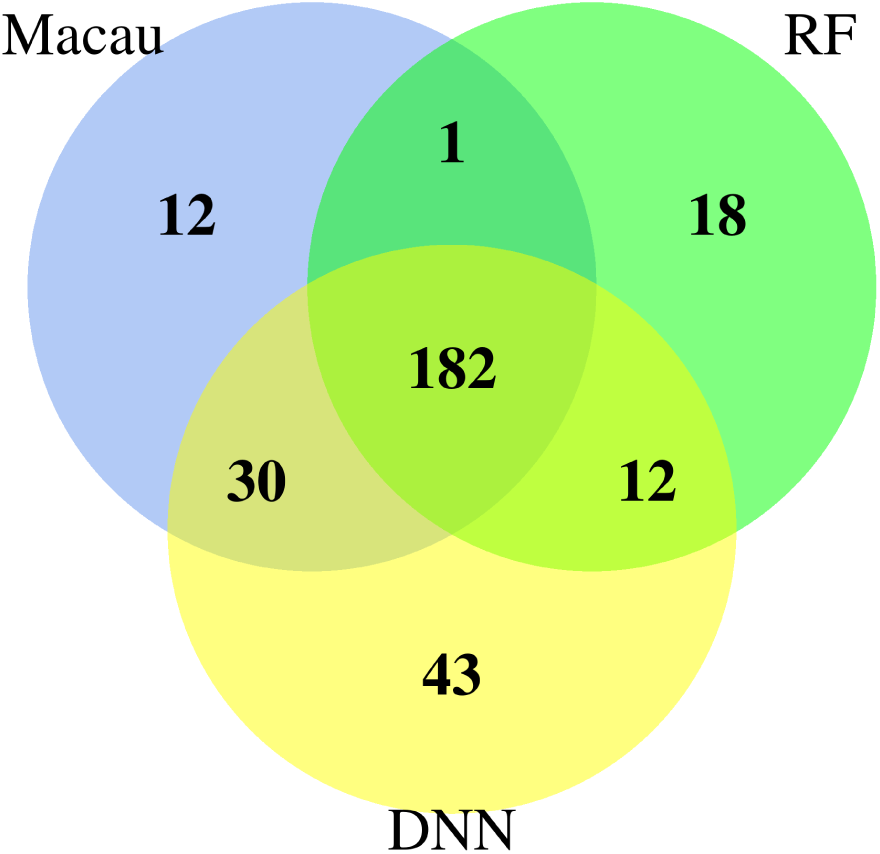
Venn diagram: number of protein targets with predictivity *AUC* > 0.7.

## References

1. Walter T, et al. (2010) Visualization of image data from cells to organisms. Nature methods 7:S26–S41.

2. Pepperkok R, Ellenberg J (2006) High-throughput fluorescence microscopy for systems biology. Nature Reviews Molecular Cell Biology 7(9):690–696.

3. Starkuviene V, Pepperkok R (2007) The potential of high-content high-throughput microscopy in drug discovery. British journal of pharmacology 152(1):62–71.

4. Yarrow J, Feng Y, Perlman Z, Kirchhausen T, Mitchison T (2003) Phenotypic screening of small molecule libraries by high throughput cell imaging. Combinatorial chemistry & high throughput screening 6(4):279–286.

5. Held M, et al. (2010) Cellcognition: time-resolved phenotype annotation in high-throughput live cell imaging. Nature methods 7(9):747–754.

6. Singh S, Carpenter AE, Genovesio A (2014) Increasing the content of high-content screening an overview. Journal of biomolecular screening 19(5):640–650.

7. Carpenter AE, et al. (2006) Cellprofiler: image analysis software for identifying and quantifying cell phenotypes. Genome biology 7(10):R100.

8. Wawer MJ, et al. (2014) Toward performance-diverse small-molecule libraries for cell-based phenotypic screening using multiplexed high-dimensional profiling. Proceedings of the National Academy of Sciences 111(30):10911–10916.

9. Caicedo JC, Singh S, Carpenter AE (2016) Applications in image-based profiling of perturbations. Current opinion in biotechnology 39:134–142.

10. Evensen L, Link W, B Lorens J (2010) Imaged-based high-throughput screening for anti-angiogenic drug discovery. Current pharmaceutical design 16(35):3958–3963.

11. Ansbro MR, Shukla S, Ambudkar SV, Yuspa SH, Li L (2013) Screening compounds with a novel high-throughput abcb1-mediated efflux assay identifies drugs with known therapeutic targets at risk for multidrug resistance interference. PLoS One 8(4):e60334.

12. Gustafsdottir SM, et al. (2013) Multiplex cytological profiling assay to measure diverse cellular states. PloS one 8(12):e80999.

13. Mayr A, Klambauer G, Unterthiner T, Hochreiter S (2016) Deeptox: toxicity prediction using deep learning. Frontiers in Environmental Science 3: 80.

14. Rogers D, Hahn M (2010) Extended-connectivity fingerprints. Journal of chemical information and modeling 50(5):742–754.

15. Baell JB, Holloway GA (2010) New substructure filters for removal of pan assay interference compounds (pains) from screening libraries and for their exclusion in bioassays. Journal of medicinal chemistry 53(7):2719–2740.

16. Bemis GW, Murcko MA (1996) The properties of known drugs. 1. molecular frameworks. Journal of medicinal chemistry 39(15):2887–2893.

17. Simm J, et al. (2015) Macau: scalable bayesian multi-relational factorization with side information using mcmc. arXiv preprint arXiv:1509.04610.

18. Breiman L (2001) Random forests. Machine learning 45(1):5–32.

## References

[1] Mayr, A., Klambauer, G., Unterthiner, T. & Hochreiter, S. DeepTox: Toxicity Prediction using Deep Learning. Frontiers in Environmental Science 3 (2016).

[2] Simm, J. et al. Macau: Scalable bayesian multi-relational factorization with side information using MCMC. arXiv preprint arXiv:1509.04610 (2015).

[3] Baell, J. B. & Holloway, G. A. New substructure filters for removal of pan assay interference compounds (PAINS) from screening libraries and for their exclusion in bioassays. J Med Chem 53, 2719–2740 (2010).

[4] Egan, W. J., Merz, K. M. & Baldwin, J. J. Prediction of drug absorption using multivariate statistics. Journal of medicinal chemistry 43, 3867–3877 (2000).

[5] Wang, Y. et al. In silico adme/t modelling for rational drug design. Quarterly reviews of biophysics 48, 488–515 (2015).

[6] Garg, P. & Verma, J. In silico prediction of blood brain barrier permeability: an artificial neural network model. Journal of chemical information and modeling 46, 289–297 (2006).

[7] Cireşan, D. C., Meier, U., Gambardella, L. M. & Schmidhuber, J. Deep Big Multilayer Perceptrons for Digit Recognition. In Montavon, G., Orr, G. B. & Muüller, K.-R. (eds.) Neural Networks: Tricks of the Trade, 581–598 (Springer, 2012).

[8] Srivastava, N., Hinton, G., Krizhevsky, A., Sutskever, I. & Salakhutdinov, R. Dropout: A Simple Way to Prevent Neural Networks from Overfitting. Journal of Machine Learning Research 15, 1929–1958 (2014).

[9] Caruana, R. Multitask Learning. Machine Learning 28, 41–75 (1997).

[10] Baumann, D. & Baumann, K. Reliable estimation of prediction errors for qsar models under model uncertainty using double cross-validation. Journal of cheminformatics 6, 1 (2014).

[11] Hochreiter, S. & Obermayer, K. Gene selection for microarray data. Kernel methods in computational biology 319 (2004).

[12] Breiman, L. Random forests. Machine learning 45, 5–32 (2001).

[13] Polishchuk, P. G. et al. Application of random forest approach to qsar prediction of aquatic toxicity. Journal of chemical information and modeling 49, 2481–2488 (2009).

[14] Oshiro, T. M., Perez, P. S. & Baranauskas, J.A. How many trees in a random forest? In International Workshop on Machine Learning and Data Mining in Pattern Recognition, 154–168 (Springer, 2012).

[15] Louppe, G. Understanding Random Forests: From Theory to Practice. arXiv preprint arXiv:1407.7502 (2014).

[16] Wright, M. N. & Ziegler, A. ranger: A fast implementation of random forests for high dimensional data in C++ and R. arXiv preprint arXiv:1508.04409 (2015).

